# Grouping cells in primate visual cortex

**DOI:** 10.1101/2024.01.16.575953

**Authors:** Tom P. Franken, John H. Reynolds

## Abstract

Our perception of how objects are laid out in visual scenes is remarkably stable, despite rapid shifts in the patterns of light that fall on the retina with each saccade. One mechanism that may help establish perceptual stability is border ownership assignment. Studies in macaque area V2 have identified border ownership neurons that signal which side of a border belongs to a foreground surface. This signal persists for hundreds of milliseconds after border ownership has been rendered ambiguous by deleting the stimulus features that distinguish foreground from background. Remarkably, this signal survives eye movements: border ownership neurons also exhibit border ownership signals *de novo* when an eye movement places the newly ambiguous border within their receptive field. The grouping cell hypothesis proposes the existence of hypothetical grouping cells in a downstream brain area. These cells would compute persistent proto-object representations and therefore have the properties to endow cells in upstream brain areas with selectivity for border ownership. Such grouping cells have been predicted to show a centripetal and persistent pattern of preferred side of ownership for a border placed parallel to the perimeter of their classical receptive field, and such a centripetal ownership preference pattern should also occur *de novo* in these same cells if an ambiguous border lands in their receptive field after a saccade. It is unknown if grouping cells exist. Here we used laminar multielectrodes in area V4 – the main source of feedback to V2 – of behaving macaques to determine whether such grouping cells exist. Consistent with the model prediction we find a substantial population of neurons with these properties, in all laminar compartments, and they exhibit a response latency that is short enough to act as the source that endows neurons in V2 with selectivity for border ownership. While grouping cell activity provides information about the location of foreground surfaces, these neurons are, counterintuitively, not as strongly tuned for luminance contrast polarity, a feature of those surfaces, as are border ownership cells. Our data suggest a division of labor in which these newly discovered grouping cells provide spatiotemporal continuity of segmented surfaces whereas border ownership cells link this location information with surface features such as luminance contrast.

## INTRODUCTION

A fundamental task for vision is to parse incoming sensory information into an organized collection of objects, so that we can efficiently interact with our environment ^1,2^. A key step in this computation is to determine, for each border between two neighboring regions in a visual scene, which side is foreground. This results in border ownership: foreground objects are perceived to “own” their borders ^3^. Rüdiger von der Heydt and colleagues discovered that 30-50% of neurons in extrastriate visual areas V2 and V4 in the primate brain are selective for border ownership ^4^. These border ownership neurons respond differently to a border in their classical receptive field (cRF) depending on whether it is owned by one side or the other side, even if the stimulus cues that define the side of foreground occur far outside of the cRF ^4–7^. For example, the blue border ownership cells illustrated in **Figure 1A** respond strongly to a vertical border in their cRF if it is owned by a figure on the right (thus B_1R_ is active in the example scene), but not if is owned by a figure on the left (thus B_2R_ is not active in the example scene). The opposite is true for border ownership cells B_1L_ and B_2L_. Neural border ownership is computed within tens of milliseconds after response onset, consistent with the notion that these signals drive visual scene segmentation ^3,4,7,8^.

**Figure 1.**
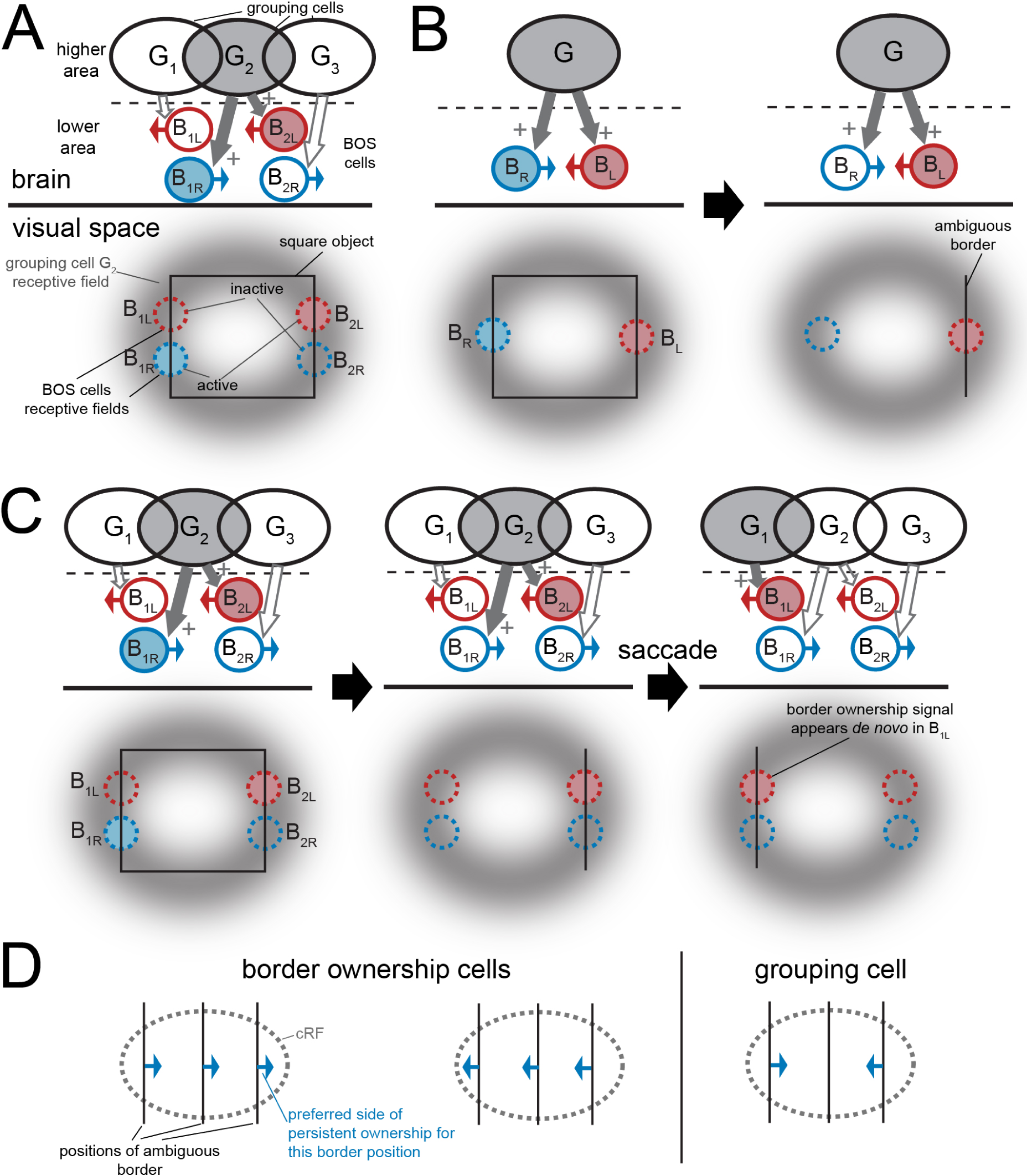
Grouping cell hypothesis. (A) Grouping circuit architecture. Border ownership (BOS) cells are labeled with B, grouping cells are labeled with G. Arrows on border ownership cells indicate their preferred side of ownership. Filled symbols indicate which neurons are active when a square object is positioned as indicated below the horizontal line. Feedback from grouping cell G_2_ endows cells B_2L_ and B_1R_ with border ownership selectivity directed towards the center of the cRF of neuron G_2_. Border ownership cells B_1L_ and B_2R_ are not strongly responsive to this square object because they do not receive excitatory feedback from G_2_, but instead from (respectively) grouping cells G_1_ and G_3_. (B) Grouping cell activity persists when the object’s borders are removed except for one (right), resulting in persistent border ownership selectivity of cell BL in this example. (C) After a saccade moves the ambiguous border over the visual scene, the persistent activity gets transferred at the grouping cell level (from G_2_ to G_1_), resulting in the activation of border ownership neuron B_1L_, whose preference is consistent with the persistent ownership of the remaining border, even though this neuron was not active while the square object was present (left: B_2L_ and B_1R_ responded to the square object, but not B_1L_). (D) For border ownership cells the preferred side of ownership for a border in the cRF is invariant for border position, but for grouping cells the preferred side should change with border position in the cRF in such a way that it points towards the center of the cRF (centripetal pattern).

A key open question is how border ownership signals are computed. Several lines of evidence support the hypothesis that they rely on contextual information supplied by cortico-cortical feedback ^9–12^. First, laminar studies found that border ownership stimuli are processed in deep cortical layers prior to granular input layers. A study in area V4 found a shorter latency of border ownership signals in deep layers compared to the granular (input) layer, as opposed to contrast polarity selectivity for the same scenes, or to responses to small stimuli in the cRF ^7^. In addition, recent data from area V1 indicate that activity in layers 5/6 leads that in layers 4C and 4A/B during the processing of border ownership scenes compared to small stimuli within the cRF ^13^. Neurons in deep layers receive cortico-cortical feedback through terminals arriving in deep layers ^14^, but also often have tall apical dendrites which would allow them to sample the feedback afferents that arrive in superficial layers ^15^. Second, contextual information that modulates the firing of border ownership neurons can occur far from their cRF and this modulation begins too soon after stimulus onset to be explained by signals propagating through the slower unmyelinated intra-areal horizontal fibers, another major contextual pathway in the cortex ^8^. A third argument concerns border ownership persistence. Border ownership signals persist if all object-defining stimulus features are deleted except for the border in the cRF (the remaining border is at that point thus ambiguous for border ownership) ^16^. This persistence of border ownership is illustrated in the lower half of **Figure 1B**: border ownership cell B_L_ responds strongly when a vertical border in its cRF is owned by an object on the left, and this activity persists if all other object borders are removed (lower right in **Figure 1B**). Moreover, this persistent border ownership signal is transferred to other neurons with eye movements, even if those neurons never responded to the object while it was still present (**Figure 1C**) ^17^. Cell B_1L_ in **Figure 1C** prefers leftward border ownership, thus it is not active when a vertical border in its receptive field is owned by a square on the right (left panel). However, when the right vertical border of this square has become ambiguous (middle panel) and then, after a saccade, lands on the cRF of B_1L_ (right panel), the persistent border ownership signal appears *de novo* in this neuron. These findings suggest that a persistent object location signal exists in a higher brain area, which provides the contextual information necessary to compute border ownership.

It has been proposed that this signal is implemented by (hypothetical) grouping cells (labeled G in **Figure 1A-C**) ^9^. Whereas the border ownership cells in **Figure 1** only ‘see’ a portion of one of the object borders in their cRF, the grouping cells, which are hypothesized to be downstream of the border ownership cells that they project to (and therefore have larger cRFs than the latter), could integrate multiple borders for the same object in their cRF. Excitatory feedback from a grouping cell to a neuron in an upstream area whose cRF is centered on one of these border fragments would endow that neuron with selectivity for border ownership. This is illustrated in **Figure 1A**, where grouping cell G_2_ provides feedback to cells B_1R_ and B_2L_. The feedback would enhance the activity of these upstream neurons more if the border in their cRF is part of an object centered at a position near the grouping cell’s cRF center, as compared to if it is part of an object centered farther away from the grouping cell’s cRF center. Computational models showed that this circuit can effectively compute border ownership for objects of various shapes, both in simple, artificial scenes and in complex natural scenes ^9,18^.

Despite more than twenty years of debate on the origins of neural border ownership signals, it remains unknown whether grouping cells exist ^19–21^. The neurophysiological properties of border ownership help define the properties grouping cells would need to have, giving us some clues about what neurons to look for. First, the border ownership computation is fast: border ownership neurons respond differently to stimuli with opposite border ownership within ∼30 msec after response onset ^4,7^. Grouping cells must thus both (1) quickly signal object location and (2) be located close enough to border ownership cells to rapidly provide the grouping signals needed to compute border ownership within this 30 msec window. The most upstream area with a large population of border ownership neurons is area V2 ^4^. Anatomically, the dominant source of feedback to V2 is area V4 ^22–25^, motivating the possibility that V4 might contain grouping cells. Second, there is reason to believe that the persistent border ownership signal that occurs after the stimulus is rendered ambiguous is not held by the border ownership cells themselves. The persistent border ownership signal disappears when the border is removed from the border ownership neuron’s receptive field, i.e. firing of border ownership neurons returns to the same baseline rate regardless of which side of the border was figure and which was ground. Surprisingly, the border ownership signal then reappears when, after a brief delay, the ambiguous border is returned to the receptive field, even though the scene does not contain information anymore about the side of ownership ^16^. Because the border ownership signal was eliminated during the pause, this suggests that the memory that enabled the border ownership signal to return resides in neurons other than the border ownership neurons themselves, i.e. in the (hypothetical) grouping cells ^16,20^. Grouping cells are thus predicted to show persistent firing to a border if that border was recently owned by an object on the side of the border towards the center of their cRF (a “centripetal” pattern of persistent ownership preference) ^16^ (**Figure 1B**, top right; **Figure 1D**, right). Third, grouping cells should also exhibit this persistent centripetal preference pattern after a saccade brings an ambiguous border in their cRF. This prediction stems from the observation that, as described above, persistent border ownership signals transfer from one population of border ownership neurons to another with eye movements if an ambiguous border lands in the cRF of the later population of (newly stimulated) neurons (even if those neurons never responded to the object before eye movement onset) ^17^ (**Figure 1C**). The centripetal preference pattern stands in contrast to the properties of border ownership neurons, whose preferred side of ownership is invariant to the position of the border in the cRF (**Figure 1D**). Armed with these predicted characteristics of grouping cells, we employed laminar multielectrode probes in area V4 of behaving macaques to determine whether this area contains neurons that exhibit the neurophysiological signatures of grouping cells.

## RESULTS

The basic stimulus paradigm is illustrated by the stimulus panels in **Figure 2A**. After the animal fixated a small target on a blank screen, we introduced an isoluminant light (or dark) square on an isoluminant dark (or light) background (left panels in **Figure 2A**). After 500 ms, this scene was replaced with a scene in which only one of the square’s borders remained and this border was prolonged over the screen (right panels in **Figure 2A**). After the display switch the ownership of this border has thus become ambiguous in the scene, since no stimulus information is present that indicates which side of the border is foreground. Indeed, the stimulus panels in the top and bottom rows in **Figure 2A** illustrate that an identical ambiguous border can be obtained by starting from different square scenes. After another 1000 ms, the animal was rewarded for maintaining fixation on the fixation target throughout the trial. The square objects were of similar size (side length 3.5-5 degrees of visual angle [dva]), luminance contrast, and used in trial sequences with similar timing as in the experiments in V2 that found persistent border ownership signals (**Figure 1B**) ^16,17^. As discussed in Introduction, grouping cells in a higher area that underlie this border ownership selectivity are predicted to respond to these ambiguous borders with a specific pattern, depending on stimulus history: such cells should fire more to the ambiguous border if it used to be owned by the side centripetal relative to the center of their cRF (**Figure 1B,D**).

**Figure 2.**
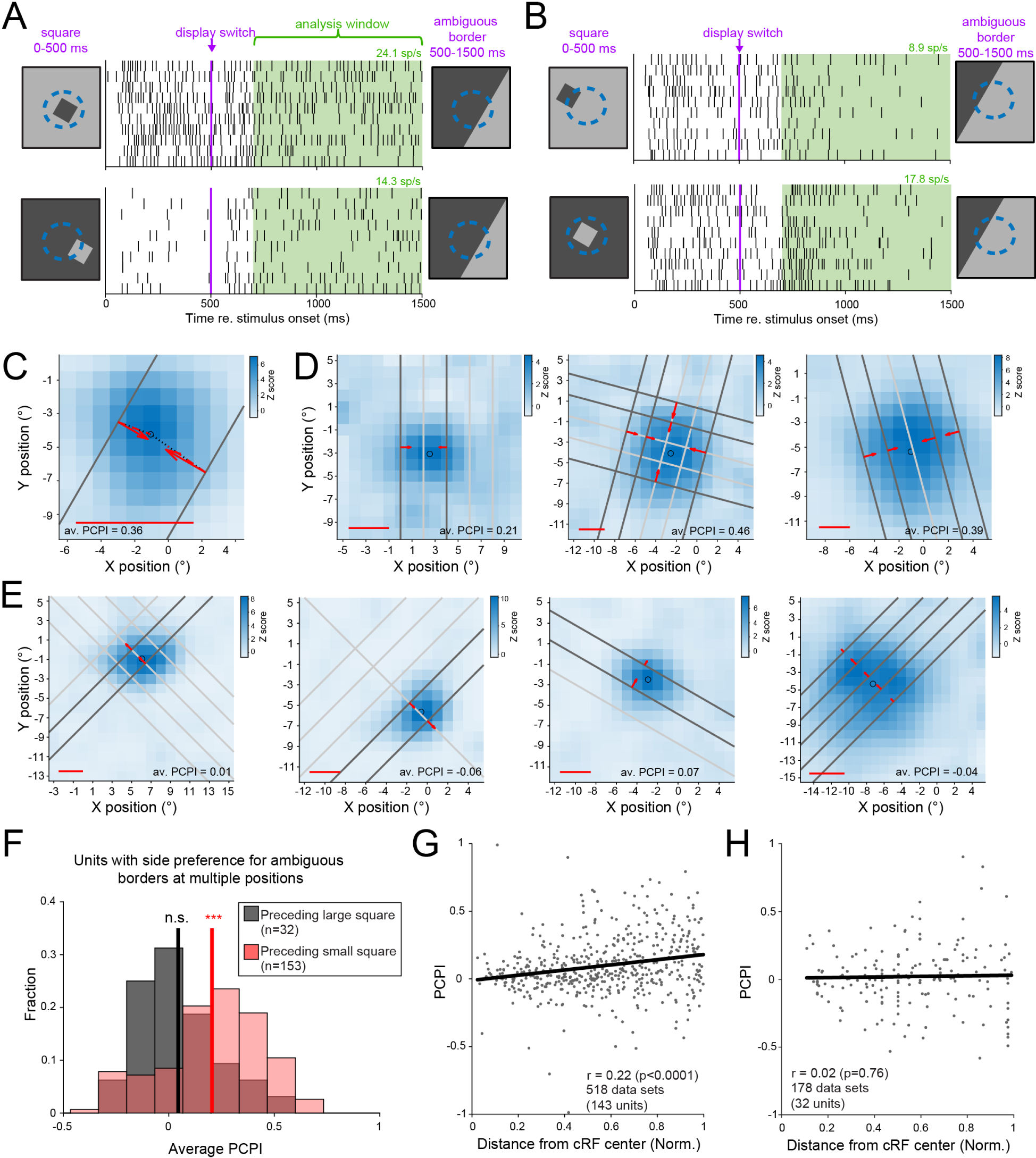
V4 responses to ambiguous borders that used to be owned by a small object show a significant persistent centripetal ownership preference. (A) Dot rasters of responses to an ambiguous luminance contrast border (500-1500 ms) that was preceded by a small square (0-500 ms), for an example single unit in V4. Green rectangle indicates the analysis window, and spike rate during this window is shown on the top right above each panel. The ambiguous border is the same for both panels, but it was preceded by a square on opposite sides of the border. Blue dashed circle indicates the cRF. (B) Similar as A, for a different position of the ambiguous border, for the same unit as in A. (C) Summary plot of the persistent side preference for the data in A,B. Grey lines show the positions of the ambiguous borders relative to the cRF. Side-preference vectors (red) show whether the response to each ambiguous border position was higher if that border was preceded by a square on one or the other side of the border (directed towards the preferred side; arrow length proportional to modulation index (MI)). Vector length projected on the line connecting the cRF center with the center of the border (black dotted line) yields the PCPI (persistent centripetal index; red dashed line). Red line below indicates length of red dotted line if PCPI would be 1. Average PCPI across both positions = 0.36. (D) Similar as B, for three other V4 units. Light grey lines indicate ambiguous border positions for which the side preference was not significant (no side-preference vector is shown for those positions). Left unit: animal Z. Middle and right unit: animal D. (E) Similar as D, for four units tested with ambiguous borders preceded by a large square for which only a portion of one of the borders fits within the cRF. Leftmost unit: animal Z; other units: animal D. (F) Histogram of PCPI for ambiguous borders that are preceded by large squares or small squares, for all single units that showed a significant side preference for multiple ambiguous border positions. The PCPI is averaged per unit across positions. Vertical lines indicate means. The average PCPI for ambiguous borders preceded by small squares is significantly larger than 0 (mean ± s.e.m. 0.20 ± 0.02, Wilcoxon signed rank test p=9.9 x 10^-16^, n = 153), as opposed to those preceded by large squares (mean ± s.e.m. 0.04 ± 0.03, Wilcoxon signed rank test p=0.4, n = 32). The difference between the distributions is statistically significant (Wilcoxon rank sum test p=1.8 x 10^-4^). (G) PCPI for different positions of the ambiguous border in the population of grouping cells. Ambiguous border position is expressed as a fraction of the radius of the cRF. Black line indicates linear fit. Slope 0.91, Pearson’s r = 0.22, < 0.0001. (H) Similar as G, for the ambiguous border positions evaluated in border ownership data sets (as in panel E; linear fit slope 0.02, Pearson’s r = 0.02, p=0.76).

### V4 contains neurons with a centripetal preference of persistent ownership for borders of small objects

Figure 2A shows the response of a V4 neuron that has the predicted property. It fires significantly more to an identical ambiguous border that is positioned in the right half of the cRF (right stimulus panels) if that border used to be owned by a small object on the left of the border (top panel) compared to when it used to be owned by a small object on the right of that border (bottom panel). To avoid including spikes that are directly evoked by the square object prior to the display switch, we limited our analysis to a window starting 200 ms after the display switch (green), similar as in the studies on persistent border ownership ^16,17^. The preferred condition for this neuron was the same irrespective of luminance contrast polarity, i.e. if we presented scenes in which the light and dark luminances were swapped in all scenes (**Figure S1A**). We computed the magnitude of the response difference as a modulation index (MI; difference in spike rates divided by sum of spike rates) across luminance conditions, equal to 0.39 for this example (significantly larger than 0, permutation test p<0.001). When we positioned the ambiguous border in the left half of the cRF for the same neuron (Figure 2B), we observed that the preferred side flips: instead of preferring a sequence in which the ambiguous border used to be owned by a square on the left, now the neuron fires more to the ambiguous border if it used to be owned by a square on the right (Figure 2B, **Figure S1B;** MI = 0.29; permutation test p=0.001). Note that this behavior is in contrast to that of border ownership neurons, who are known to prefer the same side of ownership irrespective of where the border is presented in their cRF ^4^, but instead matches the centripetal preference of persistent ownership that has been predicted for grouping cells ^20^ (Figure 1D). We summarize the physiology of this neuron in Figure 2C. The positions of the ambiguous border that were evaluated in Figures 2A**,B** are shown as dark grey lines in Figure 2C. The red side-preference vectors indicate the preferred ownership history (direction) and |MI| (magnitude) for each ambiguous border position for this neuron. Both side-preference vectors point towards the center of the cRF, as predicted for grouping cells (Figure 1D). We quantify this preference by computing a persistent centripetal index (PCPI). We project each side-preference vector on a line connecting the middle of the persistent border of the square with the center of the cRF (dotted black lines in Figure 2C). The PCPI is defined as the length of each of those projected side-preference vectors (dashed red lines in Figure 2C), and signed according to whether they are directed towards (positive) or away (negative) from the center of the cRF. We then compute the average PCPI across border positions (for the neuron in Figure 2A-C, the average PCPI is +0.36). Figure 2D shows three other example neurons that have a positive average PCPI. Light grey lines are border positions for which there was no significant preference for ownership history towards either side (permutation test p>0.05). Note that the neuron in the middle panel was tested with borders at two orientations. The population of V4 units showed a distribution of average PCPI that was significantly biased towards positive values (Figure 2F, red histogram; mean ± s.e.m. 0.20 ± 0.02, Wilcoxon signed rank test p=9.9 x 10^-16^, n = 153 units). This pattern was observed in the data from each animal (animal D: mean ± s.e.m. 0.23 ± 0.02, Wilcoxon signed rank test p=9.9 x 10^-13^, n = 111; animal Z: mean ± s.e.m. 0.13 ± 0.03, Wilcoxon signed rank test p=1.8 x 10^-4^, n = 42). This shows that the centripetal pattern of preference for persistent ownership during the ambiguous phase was significant in the population of V4 units.

To determine whether this centripetal preference is specific for objects whose borders can be grouped by V4 units (i.e. fit within the cRF), we also computed the PCPI for ambiguous borders that were preceded by large squares (side length ≥ 12 dva, excluding data sets for which the square’s corners or other sides overlapped with the cRFs (Methods)). Under these conditions, as in prior studies ^4,7^, we can define border ownership selective neurons as units that respond differently to an identical border in the cRF, depending on whether the border is owned by an object on one or the other side of the border, even though the stimulus information that defines the ownership occurs outside of the cRF. Figure 2E shows four example neurons. Now the side-preference vectors are directed to the same side of the border irrespective of their position in the cRF, as predicted for border ownership neurons (Figure 1D). This confirms that the preferred side of border ownership neurons in V4 is invariant to the position of the border (as suggested by example neurons in V2 and V4 reported by Zhou et al. ^4^), and that border ownership selectivity persists in V4 (as has been reported in V2 ^16^). For ambiguous borders preceded by large squares, the average PCPI across positions is not significantly different from 0 (Figure 2F, grey histogram; mean ± s.e.m. 0.04 ± 0.03, Wilcoxon signed rank test p=0.4, n = 32 units), and the difference between both distributions is significant (Wilcoxon rank sum test p=1.8 x 10^-4^; separately for each animal: animal D: Wilcoxon rank sum test p=0.001; animal Z: p=0.013). The centripetal pattern of persistent preference is thus specific for objects that are small enough to be grouped by the V4 cRFs, as predicted by the grouping cell model ^20^.

These data allow us to define grouping cells as units with a persistent centripetal ownership preference for ambiguous border segments, in the interval between 200 and 1000 ms after display switch to the ambiguous border, across all border positions that were evaluated for that neuron. In the following sections we will evaluate whether this subpopulation of units displays the other properties that have been predicted to occur in grouping cells.

### Grouping cells occur in all laminar compartments

Grouping cells in area V4 are hypothesized to provide the feedback signal that allows neurons in area V2 to compute the side of border ownership. This predicts that such cells are present in deep layers, in which the majority of cells that project from V4 to V2 are located ^26^. As in prior studies we used current source density (CSD) analysis to estimate laminar compartments in our recordings ^7,27–31^ (Methods). Briefly, we recorded responses to small stimuli flashed in the cRF, and computed the CSD as the second spatial derivative of the evoked local field potentials across the electrode contacts on the probe. The current sink with the shortest latency is known to correspond to the input (granular) layer, and therefore we can locate each electrode contact to superficial, input or deep layers. We find grouping cells in all three compartments (**Table**).

**Table.** Laminar distribution of grouping cells. Non-grouping cells are units that pass all inclusion criteria for grouping cells except that there is no significant centripetal pattern of ownership preference for ambiguous borders (Methods).

|  | Grouping cells (*N*) | Non-grouping cells (*N*) |
| --- | --- | --- |
| Superficial layers | 29 | 131 |
| Input layers | 56 | 146 |
| Deep layers | 49 | 174 |

Thus grouping cells indeed also occur in the deep layers from which most feedback originates.

### Grouping cells exhibit centripetal ownership preference soon after stimulus onset

To establish selectivity for border ownership in lower cortical areas by providing feedback, grouping cells need to exhibit their centripetal preference of object location earlier than the latency of border ownership signals in the recipient area. Figure 3A shows the time course of responses from the population of grouping cells. The data show that these units differentiate the centripetal from the centrifugal square position very soon after stimulus onset. The difference between the functions is shown in green in Figure 3B (latency 51.5 ms, bootstrap 95% CI [50.4 52.4] ms). We also find similarly short latencies in the subpopulation of grouping cells located in deep layers (**Figure S2**), which provide the majority of feedback to lower cortical areas. Prior studies reported latencies of 60-80 ms for the border ownership signal in area V2 for scenes with square objects of similar size and luminance contrast ^4,6,32^. The grouping signal in V4 thus starts early enough to act as the contextual signal that V2 neurons rely on to compute border ownership. For comparison we show in Figure 3C**,D** similar plots for units with persistent border ownership responses in area V4. In contrast to the grouping responses, the border ownership responses diverge later relative to response onset (Figure 3C), and the half-peak of the border ownership signal is only reached at 103.1 ms (95% CI [95.4 108.6] ms; Figure 3D, green).

**Figure 3.**
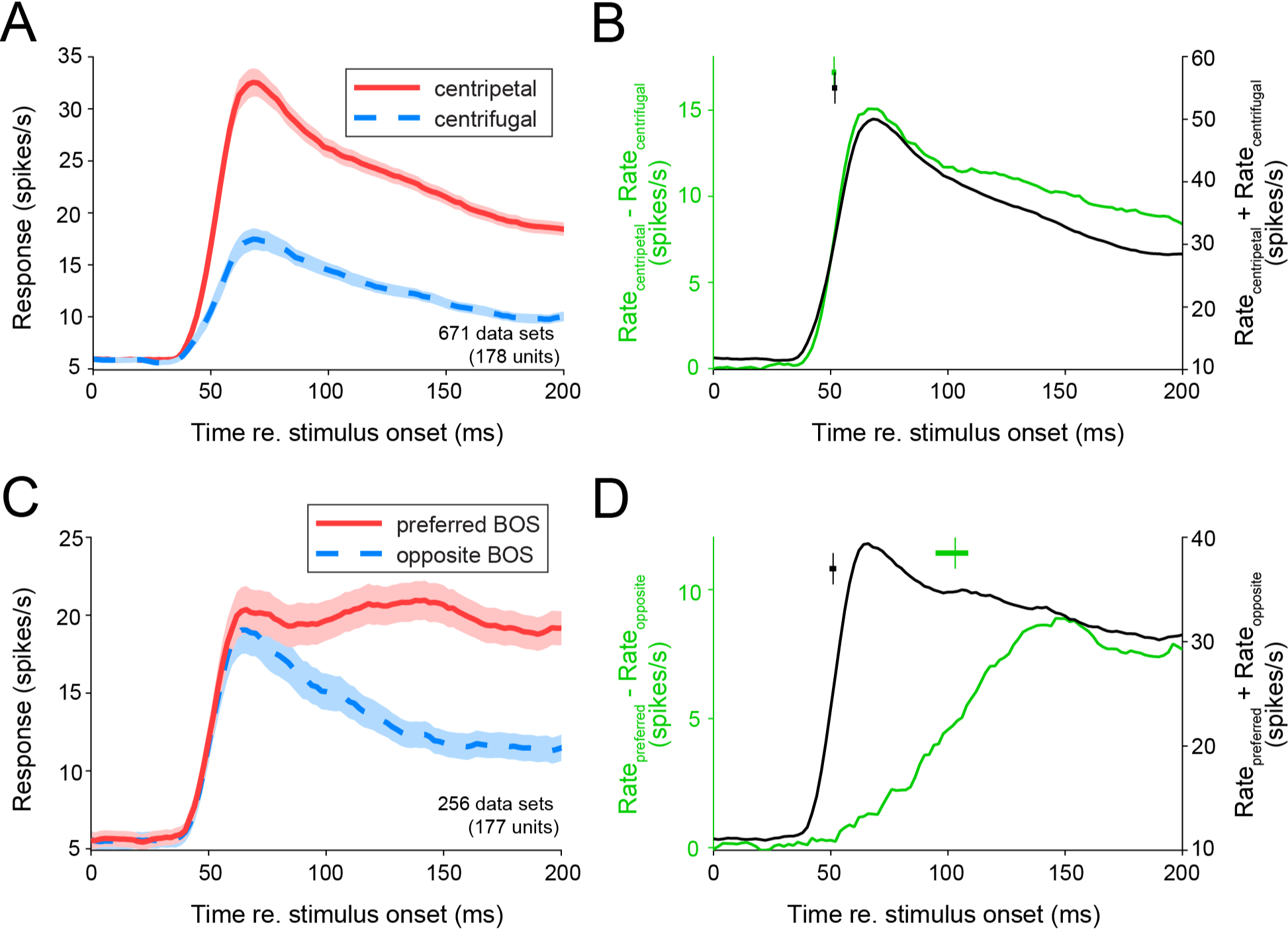
Onset of grouping signal. (A) Average time course of responses from grouping cells for centripetal (red) and centrifugal (blue dashed) square positions relative to the center of the cRF. (B) Green: difference of the response functions in A. Black: sum of the response functions in A. Symbols above functions indicate latency (difference function: 51.5 ms, bootstrap 95% CI [50.4 52.4] ms; sum function: 51.8 ms, bootstrap 95% CI [50.7 52.8] ms). (C,D) Similar as A,B, but for border ownership responses. Red and blue dashed functions in C show respectively the average response to the preferred and non-preferred side of border ownership. Latency values in D: difference function: 103.1 ms (95% CI [95.4 108.6] ms); sum function: 51.0 ms (95% CI [49.6 52.3] ms).

### Position dependence of persistent side preference in grouping cells supports fuzzy annular integration

Grouping cells are predicted to integrate object borders with a fuzzy annular template ^9^. This means that the centripetal pattern of persistent ownership preference should be more pronounced for border positions farther away from the center of the cRF (Figure 1D, right). This is indeed visible in the data from the example units in Figure 2D: the border positions closest to the center of the cRF do not show a statistically significant preference of side-of-ownership (light grey lines). To evaluate this in the population of neurons, we analyzed the relation between PCPI and the border position. We find a significantly positive correlation between the PCPI and the distance between the border and the cRF center for the population of border positions (Figure 2G; Pearson’s r = 0.22, p<0.0001). This correlation is present in the data from each animal (animal D: r = 0.30, p<0.0001, n=293; animal Z: r = 0.13, p=0.049, n=225). This relation is not predicted for border ownership selective neurons because their preferred side of ownership is invariant to the position of the border within the cRF (Figure 1D, Figure 2E). Indeed for the population of border ownership selective responses we do not find a significant correlation between border position in the cRF and persistent centripetal preference (Figure 2H; Pearson’s r = 0.02, p=0.79). Together these data support that area V4 contains grouping cells with fuzzy annular border integration templates.

### Grouping cells exhibit centripetal ownership preference for ambiguous borders after saccades

Border ownership neurons in V2 are known to signal the persistent ownership of an ambiguous border that lands in their cRF after an eye movement, even if the stimulus information that defined ownership has disappeared hundreds of milliseconds prior to response onset, and even though these neurons did not signal ownership while the object was still present ^17^. This has led to the prediction that the same population of grouping cells that support border ownership in static conditions (Figure 1A**,B**) also exhibit a centripetal pattern for persistent ownership when an eye movement positions an ambiguous border in their cRF, even if they did not signal ownership of those ambiguous borders prior to the eye movement (Figure 1C) ^20^. To test this prediction we analyzed the response of V4 single units to scene sequences that are known to result in a transfer of persistent border ownership in V2 (Figure 4A). An isoluminant square is followed by an ambiguous scene in which only one border of the square remains. After another 250 ms, a saccade is induced by moving the fixation point, and the ambiguous border only lands in the cRF after the saccade. We analyzed the spikes that occurred between 0 and 500 ms after the saccade (green in Figure 4A). We identified grouping cells independently from this task, i.e. from responses to randomly interleaved trials without saccades (square scenes followed by an ambiguous border, Figure 2). The ownership preferences of these grouping cells during the post-saccadic interval in the saccade experiment (green window in Figure 4A) are shown in Figure 4B. Each gray line indicates the position of the ambiguous border after the saccade relative to the center of the cRF, and the red side-preference vectors show the preferred side of persistent ownership for each border position. We observed that the preferred ownership vectors tend to be oriented towards the center of the cRF, resulting in a significant bias towards positive post-saccadic PCPI values (Figure 4C; mean ± s.e.m.: 0.032 ± 0.014, permutation test p=0.003, 105 data sets; animal D: 0.025 ± 0.014, p=0.03, n=90, animal Z: 0.073 ± 0.051, permutation test p=0.005, n=15). This centripetal preference was not present before the saccade (PCPI during the square stimulus for the same trials: mean −0.08 +/-SEM 0.03, permutation test p>0.999; 105 data sets). Interestingly, we found that this pattern is not a general property of V4 responses but instead is specific for grouping units: V4 units that did not pass the criterion for grouping units in the no-saccade sequences did not display a significant centripetal preference in the saccade-sequences (Figure 4D; mean ± s.e.m. 0.007 ± 0.01; permutation test p=0.14; 338 data sets). Together these data indicate that the same grouping cells that show a persistent centripetal ownership preference for objects followed by ambiguous borders also show such a pattern for ambiguous borders that land in their cRF after an eye movement, even though they did not have a preference for object location while the object was still present. This property has been predicted for grouping cells ^20^. The data support the existence of a neural population in V4 that has the properties to flexibly endow neurons in V2 with the context required to keep track of border ownership, both for changes in visual stimulation that occur during fixation or for those that are due to eye movements.

**Figure 4.**
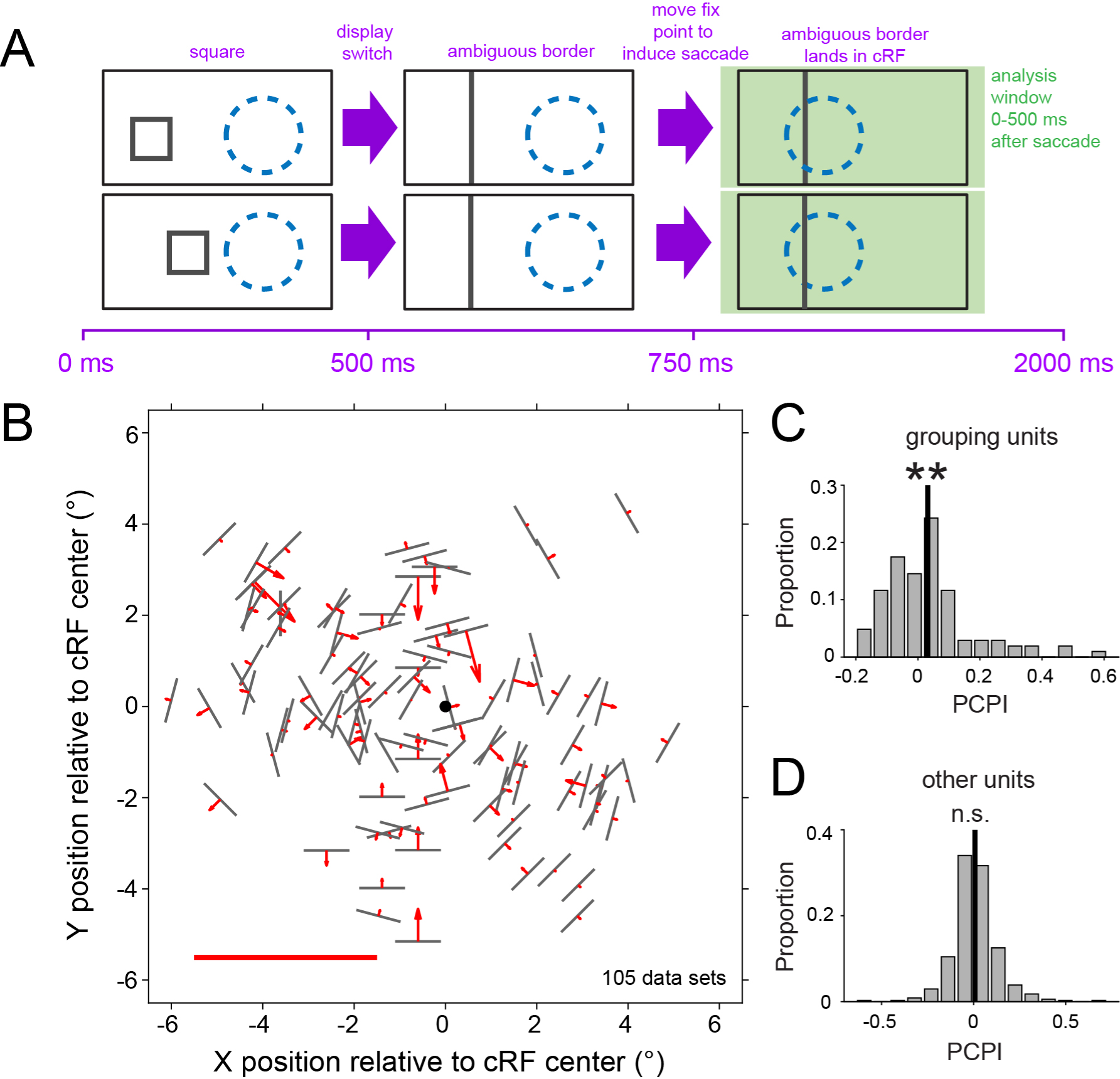
Grouping signals after saccades. (A) Sequence of events in saccade trials. Two trial sequences are shown, with the square on either side of a given border position. Squares (shown as outlines here for simplicity) were presented as isoluminant objects on an isoluminant background (similar as in Figure 2). Dashed circle represents cRF. (B) Persistent side preference after a saccade moves an ambiguous border in the cRF, for the population of grouping cells. Grouping cells were defined as units with a significant persistent centripetal preference for the same ambiguous border position in trials without saccades (see **Figure 2A,B**). Grey lines indicate for each data set (Methods) the position of the ambiguous border relative to the cRF center (black circle). Red: side-preference vectors (as in **Figure 2C**). Red line below indicates length of side-preference vector equal to 1. (C) Histogram of PCPI values for the data in B. Black line indicates mean: 0.032 (s.e.m. 0.014; permutation test p = 0.003). (D) Similar as C, for the data sets that did not pass the grouping criterion (mean ± s.e.m. 0.007 ± 0.01; permutation test p = 0.14; 338 data sets (163 units)).

### Centripetal preference of grouping units persists during interruption of the ambiguous border

When the border in the cRF becomes ambiguous after removing the stimulus cues that define the side of foreground, border ownership neurons in V2 persistently signal the side of ownership for hundreds of milliseconds (Figure 1B) ^16,17^. This persistent border ownership signal disappears when the ambiguous border in the cRF is removed, but reappears when it is reintroduced ^16^. It has therefore been suggested that the short-term memory for border ownership is not stored at the level of border ownership neurons in V2, but instead at the level of grouping cells in V4 ^20^. We therefore tested whether the grouping cells that we identified maintain their centripetal preference for the persistent ownership of ambiguous borders when the border is interrupted. We recorded responses from V4 units during similar scene sequences as those used in the V2 study (Figure 5A) ^16^. Square objects were replaced by ambiguous borders, and another 300 ms later, the scene was interrupted by a blank screen during 500 ms. After this interval, the ambiguous border was reintroduced. We randomly alternated these trials with sequences without interruption (as in Figure 2A). We identified grouping cells as above, from the responses to the uninterrupted sequences. Figure 5A shows the response of a grouping cell to the interrupted sequence. During the scene interruption, the difference in response between the ownership conditions persists (green interval). For the population of grouping cells, we find a persistent centripetal side-preference signal across the blank interval (Figure 5B**,C**). This stands in contrast to the population of units that do not pass the criterion for grouping cells (**Figure S3A,B**). These data support the hypothesis that grouping cells in V4 construct a persistent proto-object representation that underlies the re-emergence of border ownership signals in area V2 after border interruption, even though these had completely disappeared during the interruption ^16^.

**Figure 5.**
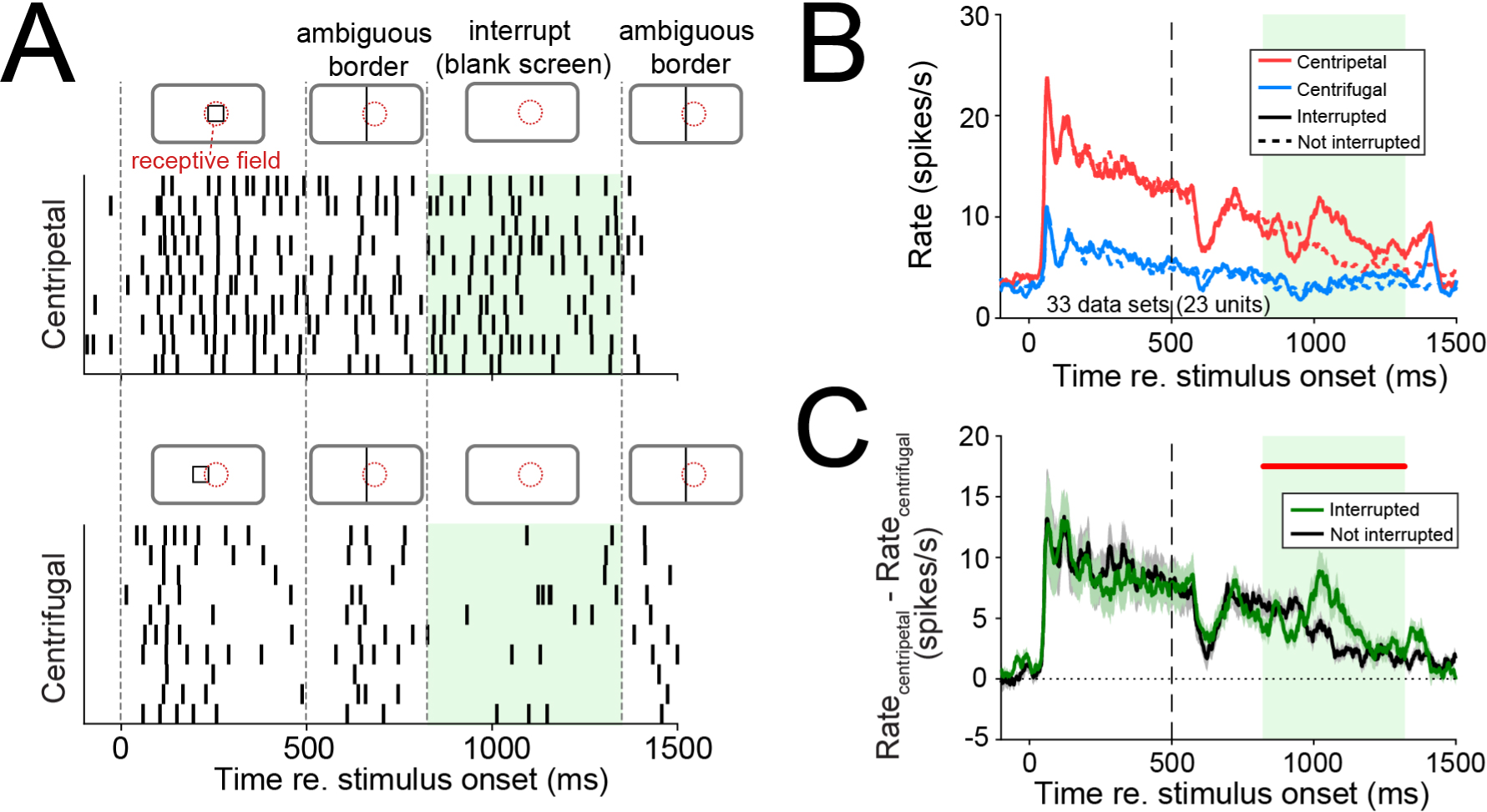
Grouping signal persists across interruption of ambiguous border. (A) Dot rasters from an example grouping cell for trials where an ambiguous border, which used to be owned by a small square object on the side of the border towards the center of the cRF (top panel) or on the opposite side (bottom panel), is temporarily interrupted (green interval: scene replaced by blank screen). Cartoons above the panels indicate stimulus sequence during the trial. For clarity the cartoons show outline figures, but squares were presented as isoluminant squares on an isoluminant background, and the square was either lighter or darker than the background (Methods). Border orientation was 150° for this data set. (B) Average time course across the population of grouping cells to trials with (solid line) or without (dashed line) interruption of the ambiguous border, plotted separately for trials where the border used to be owned by a square object towards (centripetal: red) or away (centrifugal: blue) from the center of the cRF. The experiment was done in 33 data sets from 23 grouping cells in animal D. (C) Difference in firing rate between centripetal and centrifugal functions in B (mean; shaded bands: standard error of the mean), plotted separately for trials with and without interruption of the ambiguous border. Red line indicates time points during the interruption of the ambiguous border (green rectangle) for which the spike rate difference between centripetal and centrifugal in the interrupted condition was significantly larger than 0 (permutation test p<0.05).

### Grouping cells are less selective for luminance contrast polarity than border ownership cells

It has been predicted that grouping cells would be less selective for surface features than border ownership cells ^33^. One feature that characterizes a surface is its luminance contrast relative to background. Border ownership cells are often selective for the order of luminances across a border (luminance contrast polarity) ^4,7^. An example neuron that we recorded in V4 is shown in Figure 6A. This neuron is selective for border ownership, it fires more spikes when an identical horizontal border in its cRF is owned by a square figure below it compared to when it is owned by a square figure above it, even if the stimulus information in the cRF is identical (compare 1 with 3, and 2 with 4). But the response to scenes with the same side-of-figure also differs depending on contrast polarity: the neuron fires more if the area below the border has a higher luminance than the area above the border (compare 2 with 1, and 4 with 3). We quantified the tuning for both side-of-figure and contrast polarity using the reliability metric introduced by Zhou et al. ^4^. Briefly, this metric is computed by comparing the spike counts between 10,000 randomly selected sets of four trials (in each set, one trial is randomly sampled from each of the four conditions). Reliability for side-of-figure is then defined as the proportion of such sets for which the spike count is highest for the side-of-figure that evokes the highest spike rate across trials (side-of-figure ***a*** in the example in Figure 6A). The reliability for the data in Figure 6A for side-of-figure is 1.000: comparing spike counts between such trial sets always results in higher spike counts for side-of-figure ***a*** compared to side-of-figure ***b***. We can compute a similar reliability metric for luminance contrast polarity. For this same neuron we find that reliability for luminance contrast polarity is 0.9994 (for almost all sets there is a higher spike count for luminance contrast polarity ***d*** compared to luminance contrast polarity ***c***). In the population of border ownership cells we find that high reliabilities for side-of-figure and for luminance contrast polarity often co-occur in the same units (clustering of data points in top right corner in Figure 6C). This confirms that border ownership responses in V4, as in V2, are often highly reliable for luminance contrast polarity ^4^.

**Figure 6.**
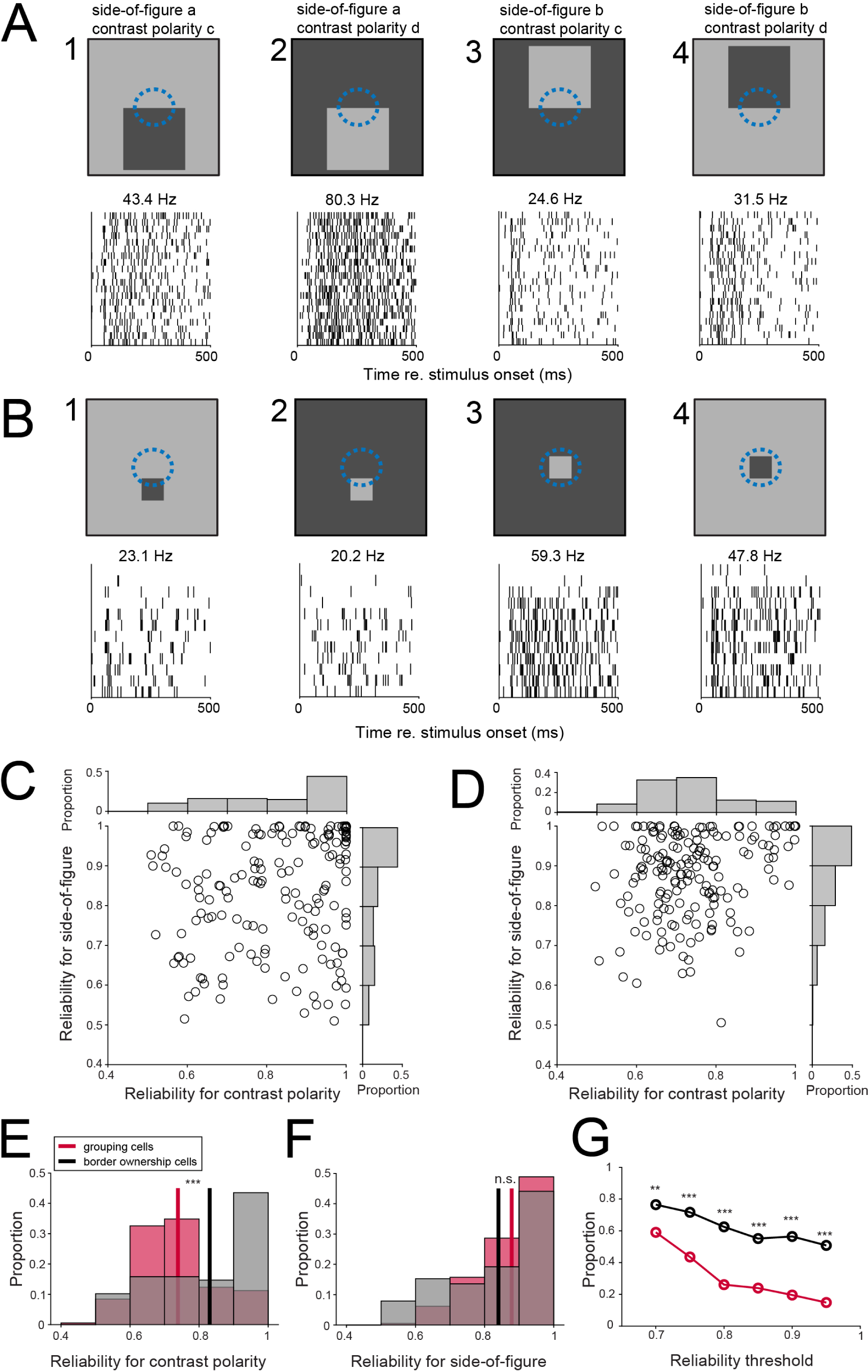
Reliability for contrast polarity and side-of-figure in grouping cells and border ownership cells. (A) Dot rasters of responses from an example border ownership cell to four scenes. Average firing rate is shown above each dot raster. (B) Similar as A, for responses from a grouping cell. (C) Reliability for luminance contrast polarity (abscissa) and side-of-figure (ordinate) are shown for the population of border ownership cells (177 units). Histograms indicate marginal distributions. Reliability was computed using spikes recorded during the square phase (0-500 ms after stimulus onset). (D) Similar as C, for the population of grouping cells (178 units). (E) Distribution of reliability for luminance contrast polarity for grouping cells and border ownership cells. Vertical lines indicate means (grouping cells: 0.74 (n=178), border ownership: 0.83 (n=177), Wilcoxon rank sum test p=2.3 x 10^-9^). (F) Similar as E, for reliability for side-of-figure (mean grouping: 0.88, mean border ownership: 0.84, Wilcoxon rank sum test p=0.10). Colors as in E. (G) Proportion of units whose reliability for luminance contrast polarity is higher than a given threshold, as a fraction of the number of units that pass the same threshold for reliability for side-of-figure. Colors as in E. **: permutation test p<0.01. ***: permutation test p<0.001.

The grouping cell in Figure 6B shows a different picture. While reliability for side-of-figure is high (0.96), reliability for contrast polarity is lower (0.71), because the spike counts between scenes with opposite contrast polarity do not differ much. This trend was confirmed at the population level (Figure 6D). Reliability for luminance contrast polarity was significantly lower for grouping cells than for border ownership cells (Figure 6E, grouping: mean 0.74, border ownership: mean 0.83, Wilcoxon rank sum test p=2.3 x 10^-9^; animal D: p=2.2 x 10^-6^ (141 grouping cells and 132 border ownership cells); animal Z: p=0.027 (37 grouping cells and 45 border ownership cells)), as opposed to reliability for side-of-figure (Figure 6F, grouping: mean grouping: 0.88, mean border ownership: 0.84, Wilcoxon rank sum test p=0.10). The co-occurrence of high reliability for side-of-figure and contrast polarity was significantly different between border ownership cells and grouping cells: for the subpopulation of units that pass a reliability threshold for side-of-figure, the proportion that passes the same reliability threshold for contrast polarity (ordinate in Figure 6G) is higher for border ownership cells than for grouping cells, irrespective of the threshold value. Together these data show that responses from grouping cells and border ownership cells have a similar degree of reliability for side-of-figure, but they differ in reliability for luminance contrast polarity (weak for grouping cells, strong for border ownership cells).

## DISCUSSION

Despite over twenty years of debate on the nature of border ownership computations in the visual cortex, the underlying mechanisms remain unclear ^4,12,19,34^. While multiple lines of evidence support a major role for contextual information fed back from a higher brain area ^7,8,17,35^, it remained unclear whether grouping cells, the hypothetical source of such feedback, exist ^9,20,21,33^. Our single unit recordings reveal that area V4 contains indeed a substantial population of units that have several of the properties that have been predicted for grouping cells. We found units that signal the persistent ownership of ambiguous borders in their cRF with a centripetal pattern of preferred ownership. These persistent ownership signals are stronger when the ambiguous border is positioned farther from the center of their cRF, consistent with an annular template of border integration. These same units also display such centripetal ownership patterns *de novo* after a saccade moves an ambiguous border in their cRF, as opposed to units that do not pass the grouping cell criterion. We find a substantial proportion of grouping cells in deep cortical layers, from which most of the feedback originates ^26^. Grouping cells signal ownership 10-30 ms earlier than the latency of border ownership signals in V2 ^4,6,32^, thus fast enough for the ∼6-ms conduction delay that has been estimated for the myelinated fibers that project from V4 to V2 ^9^. Our data are thus consistent with a circuit in which contextual signals from grouping cells in V4 endow neurons in area V2, the lowest area in the visual hierarchy with a substantial proportion of border ownership neurons ^4^, with selectivity for border ownership (Figure 1).

The annular integration template that is suggested by our data is in agreement with the characterization of V4 receptive fields in other studies. One prior study used short bars presented in fast reverse correlation sequences to map the fine structure of orientation tuning in V4 ^36^. They found that the preferred orientation for border segments at different locations in the cRF can be such that it forms an annular pattern (e.g. their Figure 3, II). This is consistent with earlier work that revealed that concentric gratings evoke particularly strong responses in V4 neurons ^37^. The shortest latency that we find for grouping cells in the input layer (**Figure S2B**) suggests that this annular integration may be computed from feedforward inputs.

The remapping property of grouping cells with saccades is reminiscent of receptive field remapping that has been reported in various brain areas. Specifically, grouping cell remapping falls in the category of post-saccadic, memory trace remapping ^38,39^. This refers to neuronal responses that occur after a saccade, to a stimulus that was briefly presented before the saccade in the future cRF (i.e. the location where the cRF lands after the saccade). Memory trace remapping has been observed in parietal cortex (lateral intraparietal area), frontal eye field, superior colliculus and MST ^40–44^, Memory trace remapping was also found in visual areas V2, V3, V3A and V4, but only when the stimulus was presented after the fixation point had already been moved, briefly before the saccade occurred ^45,46^. Nakamura and Colby explored different timings in areas V2, V3 and V3A and found no remapping when the interval between stimulus and saccade was more than 200 ms ^45^. Remapping in another V4 study was also limited, to less than 100 ms before saccade onset ^47^. The remapping that we observe in grouping cells differs in several respects. First, these units show remapping of centripetal ownership patterns for square objects that disappeared more than 250 ms before the saccade. Second, the typical paradigm used in remapping studies is to flash an isolated stimulus probe on a uniform background, whereas the remapped grouping signal is the difference in response to an identical ambiguous border in the cRF with different stimulus histories. Third, we find that the remapped signal specifically occurs in those units that also show a persistent centripetal ownership preference pattern in trials without saccades, suggesting that a single population of grouping cells contributes to a continuous spatiotemporal representation both when the visual scene changes during fixation and when eye movements shift visual information over the retina. An interesting avenue for further research is to study how grouping cells shift their cRFs when saccades occur. It has been hypothesized that the pulvinar may provide routing signals that underlie such computations ^21,48^.

Grouping cells in V4 and border ownership cells in V2 each provide information about which side of a border in their cRF is foreground. This leads to the question of why one needs these different types of neurons to segment visual scenes. Our data suggest a division of labor. On the one hand, the same population of units shows centripetal ownership patterns during the object phase (Figure 3), during the ambiguous border phase (Figure 2), when the ambiguous border is interrupted (Figure 5) and after a saccade moves an ambiguous border in the cRF (Figure 4). This suggests that grouping cells construct a persistent proto-object representation that aids segmentation when informative visual information disappears (for example during occlusion) or when the visual information shifts over the retina with eye movements. Feedback from such cells would explain why border ownership cells in V2 show persistence for border ownership when only an ambiguous border remains, and why this persistent border ownership signal reappears after it completely disappeared during border interruption ^16,17^. A role for V4 in supporting a spatiotemporal continuous representation is also supported by prior work: V4 neurons are selective for the shape of partially occluded objects ^49^, and fire more when their cRF lands on an occluded object than when it lands on an identical surface that does not occlude an object ^21^. On the other hand, our data suggest that grouping cells do not encode surface features, such as luminance contrast polarity, with the same reliability as border ownership cells. Grouping cells may thus be a subset of the feature-invariant shape-selective cells that have been reported in V4 ^50,51^. Together a picture emerges in which grouping cells underlie spatiotemporal continuity of objects, whereas border ownership cells link this location information to object features. Separate neural populations for encoding the spatiotemporal continuity of objects and their features are consistent with longstanding findings regarding apparent motion in perception. If two objects are flashed in rapid sequence at different locations, they are perceived as one object, even if the two flashed objects differ in their attributes, such as shape or color ^52–54^.

Our data lay the foundation for a causal study of the grouping cell model. This model predicts that inactivation of the V4 cells that we identified here decreases the selectivity for border ownership of V2 neurons if their preferred side of ownership corresponds to the cRF center of the inactivated grouping cells. It is possible that border ownership neurons in V4 also contribute to border ownership selectivity in V2, as prior research showed that neurons in deep layers of V4, from which cortico-cortical feedback originates, compute border ownership earlier than neurons in V2 ^4,7,32^. In addition, a comparison of the latencies of the contextual modulations evoked by object fragments in different locations in the surround suggested that (slower) intra-areal horizontal connections may help convey contextual information that is located close to the cRF ^8,34^. Our experiments revealed the first direct recordings from neurons in V4 that have several of the properties that had been predicted for grouping cells, which may underlie border ownership selectivity in early visual areas through cortico-cortical feedback.

## ACKNOWLEDGEMENTS

This work was supported in part by NIH grant number EY031795 (to T.P.F.), a postdoctoral fellowship from the George E. Hewitt Foundation for Medical Research (to T.P.F.), a NARSAD Young Investigator Grant from the Brain and Behavior Research Foundation (to T.P.F.), the Fiona and Sanjay Jha Chair in Neuroscience (to J.H.R.). We thank Dr. Mathias LeBlanc, Dr. Sean Adams, and Ms. Catherine Williams for excellent animal care.

## AUTHOR CONTRIBUTIONS

Conceptualization, T.P.F. and J.H.R; Formal analysis, T.P.F.; Funding acquisition, T.P.F. and J.H.R; Investigation, T.P.F.; Software, T.P.F.; Supervision, J.H.R.; Visualization, T.P.F.; Writing – Original Draft, T.P.F.; Writing – Review and Editing, T.P.F. and J.H.R.

## STAR METHODS

### Animals

Two male rhesus macaques (*Macaca mulatta*), aged 13 years (animal Z) and 15 years (animal D) were used in these experiments. All of these procedures conform to the Guide for the Care and Use of Laboratory Animals and were approved by the Institutional Animal Care and Use Committee at the Salk Institute for Biological Studies (protocol 14-00014).

### Surgery

A titanium recording chamber was installed over dorsal area V4 using previously published procedures ^7^. In a separate surgical session, the dura over the cortex in the chamber was removed and replaced with a transparent silicone artificial dura ^55^.

### Electrophysiology

Recording procedures have been described before ^7^. Briefly, 32-channel linear silicon multielectrode probes (100 µm electrode pitch, ATLAS Neuroengineering) were lowered through the artificial dura in the cortex with a hydraulic Microdrive (MO-972A, Narashige). Voltage signals were continuously monitored and recorded using Intan hardware (RHD2132 amplifier board and RHD2000 amplifier evaluation system, Intan Technologies LLC). Probes were advanced until multiunit activity was visible on the deepest ∼2.6 mm of the probe.

### Stimulus presentation and task control

Visual stimuli were presented using a LED projector (Propixx, VPixx Technologies) that displayed the images on a rear-projection screen (Stewart Filmscreen). Stimulus presentation and task were controlled by MonkeyLogic software (https://www.brown.edu/Research/monkeylogic/; https://monkeylogic.nimh.nih.gov/; ^56^). A photodiode was used to measure stimulus timing. Eye position was monitored and recorded using an infrared eye-tracking camera (ETL-200, ISCAN). Trials were aborted if eye position deviated from the fixation point (threshold typically 1 dva radius).

### Receptive field mapping stimuli

cRFs were mapped using the same subspace reverse correlation technique that we have used before ^7^. We used static Gaussian-windowed square wave gratings (FWHM 2 dva; 6 orientations; 2 contrast polarities; typical luminance contrast 80%; spatial frequency phase such that one contrast edge was positioned centrally in the window; one half was one of seven colors or gray scale, the other half was always grayscale) and dark gray rings (luminance contrast 80%; diameter 2 dva; ring thickness 0.25 dva) that were presented every 50-60 ms at random locations selected from a 25 dva x 25 dva grid (spacing 1 dva between adjacent grid locations) centered on the appropriate visual quadrant. To appropriately position and size the other stimuli during the recording, the aggregate cRF of the units recorded on the probe was estimated during the recording session by analyzing high-gamma filtered local field potentials.

### Orientation tuning stimuli

Similar stimuli as those for cRF mapping were used, but with a circular window, and sized and positioned such that the grating covered the estimated cRF (12 orientations; 2 contrast polarities; luminance contrast 54%; stimulus duration 200 ms). We used this data set to estimate the preferred orientation and color contrast during the recording session, by analyzing high-gamma filtered local field potentials.

### Layer assignment

To estimate the laminar position of the electrode contacts on the probe, we used the same current-source density (CSD) mapping procedure that we have used before ^7,27^. Briefly, evoked local field potentials (LFP) were recorded for dark gray ring stimuli (luminance contrast 94%, size and position such that the stimulus falls within the estimated cRF, stimulus duration 32 ms). The CSD was computed as the second spatial derivative of the LFP. In the resulting pattern of current sinks and current sources, we could identify the granular (input) layer as the current sink with the shortest latency based on previous physiological and histological studies ^7,28,29,31,57^, and thus infer which electrode contacts were positioned respectively in superficial and in deep cortical layers.

### Border ownership and grouping stimuli and task

Based on the online analysis of cRF mapping and orientation tuning stimuli, position, size, color and orientation of the squares for border ownership and grouping scenes were chosen. Similar as in prior research, an isoluminant square was positioned on an isoluminant background (luminance contrast 54% ^4,7^). Square sizes were either large (between 12 dva x 12 dva and 20 dva x 20 dva, so that only a portion of the central border would overlap with the cRF and border ownership selectivity can be defined), or small (between 3.5 dva x 3.5 dva and 5 dva x 5 dva; so that the square can be grouped within the cRF and grouping cells can be defined). Note that the ∼4 dva square size is the same as used in V2 studies to identify border ownership cells ^4,16,17^. As in those studies, square and background were either both grays, or a combination of gray and a color. In a basic set of 4 scenes, square position could be on either side of a shared border (referred to as the central border) such that the luminance contrast polarity was kept constant around that border (for example, scenes 1 and 3 in Figure 6A (large squares); scenes 1 and 3 in Figure 6B (small squares)), and in a similar pair of scenes contrast polarity was opposite to that of the first pair (scenes 2 and 4 in Figure 6A**,B**). Trials started with the appearance of a small light gray fixation target (0.2 dva x 0.2 dva, luminance contrast 80%) that was shown on a blank grey screen (luminance set at geometric mean of the luminances in the square scenes). When the animal acquired and maintained fixation at this target for 400 ms, the square scene appeared for 500 ms. Then the scene was replaced by a scene in which all borders were deleted except for the central border, which was prolonged over the screen such that it separated two isoluminant areas (ambiguous border, right panels in Figure 2A). After another 1000 ms, this scene disappeared and the animal was rewarded with juice for maintaining fixation throughout the trial. Usually a range of conditions were used in each recording session that differed in square orientation (between 1 and 6 orientations were sampled, depending on recording time) and position of the central border (between 1 and 5 per orientation, typically spaced by 2 dva). Conditions were presented pseudorandomly in blocks such that each condition was played once before repeating conditions (typically 8-10 repetitions).

In a subset of recording sessions, we randomly interleaved these trials with similar trials in which the animal had to make a saccade if the fixation point moved. In saccade trials the fixation target was initially positioned such that the cRF would not contain any of the square’s borders. The fixation target was moved to its final position 250 ms after the square had been replaced with the ambiguous border (thus similar as in a border ownership study in macaque V2 ^17^). The animal needed to saccade within 500 ms to the new position (saccade distance between 7 dva and 12 dva), after which the ambiguous border scene remained for another 1000 ms and the animal was rewarded. These saccade trials were randomly interleaved with non-saccade trials, similar as above, in which the fixation target was at the final position from the start.

In the border interruption experiment (Figure 5), the ambiguous border scene was temporarily replaced with a blank grey screen, 800 ms after square onset and for a period of 500 ms, after which the same ambiguous border scene was reintroduced.

### Analysis

Data were analyzed in MATLAB (MathWorks, Natick, MA). Statistical significance was defined as p<0.05.

### Spike sorting

The voltage data were sorted offline with SpyKING CIRCUS ^58^. The resulting clusters were curated manually using the SpyKING CIRCUS MATLAB GUI. Single units were identified based on a well-defined refractory period in the interspike interval histogram, and are referred to in the manuscript as ‘units’.

### Receptive field mapping

To measure the cRF, we computed the mean spike counts for each stimulus position in a window [30 100] ms after stimulus onset. These values were transformed to *z*-scores by subtracting the mean and then dividing by the standard deviation of spike counts in an equally long window preceding stimulus onset ^7,59^. These *z*-scores were arranged in a map of visual space according to the stimulus positions, which was then smoothed with a Gaussian filter (using the MATLAB function *imgaussfilt* with parameter σ = 1). We defined the cRF contour using the MATLAB function *contourc* at *z* = 3. The center of the cRF was defined as the centroid of this contour computed using MATLAB function *centroid*. For the analyses in Figures 2G**,H**, the distance between the center of the central border in the square scene sequences and the center of the cRF was normalized by dividing it by the radius of a circle with an identical surface area as the cRF area within contour *z* = 3.

### Inclusion criteria

In the following, a data set refers to a collection of responses to four distinct sequences that have identical central borders but that differ in side of ownership and/or luminance contrast polarity (see *Border ownership and grouping stimuli and task*). We used the following inclusion criteria for a data set to be included in the analysis: 1) at least 6 repetitions are available for each unique trial sequence; 2) the unsigned angle between a line connecting the middle of the central border and the RF center, and a line orthogonal to the central border is larger than 45° and smaller than 135° (this ensures that the preferred square position is unambiguously towards or away from the cRF center); 3) average spike rate is >1 spikes/s for at least one of the four conditions, both during the square phase (window [0 500] ms after square onset) and during the ambiguous border phase ([700 1500] ms after square onset); 4) the evoked spike count during the square phase for at least one of the four conditions, across available trials, is significantly different from that in an equally long window just prior to square onset across all conditions (two-sided Wilcoxon rank sum test with Bonferroni correction). For large square data sets (used to identify units that are selective for border ownership), an additional criterion is that the central border intersects the cRF such that the distance between any part of the cRF contour and any part of the noncentral square borders is ≥1 dva ^7^.

### Evaluation of persistent ownership

To evaluate the response differences between sequences with identical central border but different preceding square position, we analyzed spike counts evoked during the ambiguous border phase, in a window [700 1500] ms after onset of the square stimulus (i.e. starting 200 ms after the display switch from square to ambiguous border). We first computed a modulation index MI defined as the difference divided by the sum of responses to a data set:

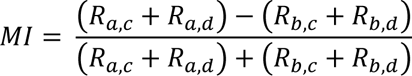

where *R*_*i,j*_ represents the average spike rate in the analysis window for square position *i* and luminance contrast polarity *j*. Note that during the analysis window the square has already disappeared, i.e. square position refers to the side of the central border that used to be owned by the square in the same trial (referred to as the ‘preceding square’). Statistical significance of |*MI|* was evaluated using a permutation test: we shuffled the ownership stimulus labels 10,000 times, separately for each value of luminance contrast polarity, and the p value was estimated as the fraction of |*MI_shuffled_*| that was at least as large as |*MI*|. Side-preference vectors (red vectors in Figure 2C-E) originate from the middle of the central border of the preceding square, have magnitude equal to |*MI*| and orientation orthogonal to the central border. Their direction is defined as the position of the preceding square relative to the border that resulted in the highest spike count during the ambiguous phase. The persistent centripetal index (PCPI) is the length of the side-preference vector projected on a line connecting the middle of the preceding square border and the center of the cRF, and the PCPI is signed according to whether this projected vector point towards the cRF center (positive) or away from the cRF center (negative). The average PCPI for a single unit is the average PCPI across all available border positions. The correlation between PCPI and distance between the middle of the central border of the preceding square and cRF center (Pearson correlation coefficient and p value) was analyzed with MATLAB function *corrcoef* (Figure 2G**,H**).

### Grouping cells and border ownership cells

To identify grouping cells, we analyzed the trial sequences where the ambiguous border was preceded by a small square. All data sets for a cell that passed the inclusion criteria were combined and the square position labels in each data set were re-labeled according to whether the preceding square position was on the same side (centripetal) or the opposite side (centrifugal) as the cRF center, compared to the central border in the data set. Spike trains were then divided in four groups, according to this preceding square position (centripetal or centrifugal) and luminance contrast polarity (2 possibilities: higher luminance either towards or away from cRF center). We then computed a modulation index *MI* similar as above, for spike counts during the ambiguous border phase ([700 1500] ms after square onset), signed such that positive means centripetal side has higher spike count, and negative means the centrifugal side has higher spike count. We assessed statistical significance by comparing |*MI*| with a null distribution created by shuffling the square position labels (10,000 shuffles). The p value was estimated as the fraction of |*MI_shuffled_*| that was at least as large as |*MI*|. We defined grouping cells as those for which the *MI* was positive (i.e. centripetal preference) and p<0.05.

To identify border ownership cells, we analyzed the trial sequences where the ambiguous border was preceded by a large square that passed all inclusion criteria. We performed a similar permutation test on |*MI*| using square position (not relabeled according to cRF position) and luminance contrast polarity. Border ownership cells were defined as those for which p<0.05.

### Response time course

Time course of grouping and border ownership signals was computed by binning spikes in 2 ms bins and then convolving the resulting vectors with a postsynaptic kernel *K*(*t*) ^60^

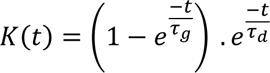

where τ_*g*_ = 1 ms and τ_*d*_ = 20 ms. The resulting functions were averaged per condition, and then across both contrast polarities for each value of border ownership or centripetal/centrifugal preceding square position. Latencies for sum and difference functions (**Figs. 3C,D**) were defined as the time point when the function crossed the value halfway between the baseline (defined as the average value during the first 10 ms) and the peak, estimated using linear interpolation (MATLAB function *interp1*). 95% confidence intervals on latencies were computed using a bootstrap analysis (MATLAB function *bootci* with 1000 bootstraps).

### Reliability analysis

We used the reliability metric introduced by Zhou et al. ^4^ to analyze the trial-to-trial reliability of selectivity for side-of-figure and contrast polarity (Figure 6), similar as in our prior work ^7^. For this analysis, spikes were counted in each trial during the square phase ([0 500] ms after square onset). For each data set, 10,000 sample sets were generated that each consisted of one trial from each of the four conditions (two side-of-figure x two contrast polarity). Each sample set thus consists of four spike count numbers. Reliability R for a given data set is then defined as

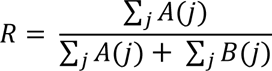

where j corresponds to the index of sample sets. *A*(*j*) and *B*(*j*) indicate whether the sign of the spike count difference between alternative values of the variable of interest for spike train set *j* is, respectively, the same or opposite compared to that of the difference between the average spike counts across all trials in the data set, for that same variable:

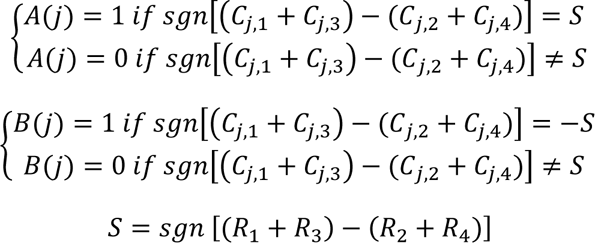

where *C*_*j*,*i*_ represents the spike count for condition *i* in spike train set *j*, *R*_*i*_ is the average spike count (across trials) for condition *i*, and *sgn* is the sign function. Conditions *i* = 1 and *i* = 3 share the same value for the variable of interest, and the same is true for conditions *i* = 2 and *i* = 4. This reliability analysis was done both for contrast polarity (abscissa in **Figs. 6C,6D**) and side-of-figure (ordinate in **Figs. 6C,6D**). Reliability was averaged across the available data sets per unit.

## DATA AVAILABILITY

MATLAB figures with embedded data will be made publicly available on figshare upon acceptance for publication.

---

**Figure S1.**
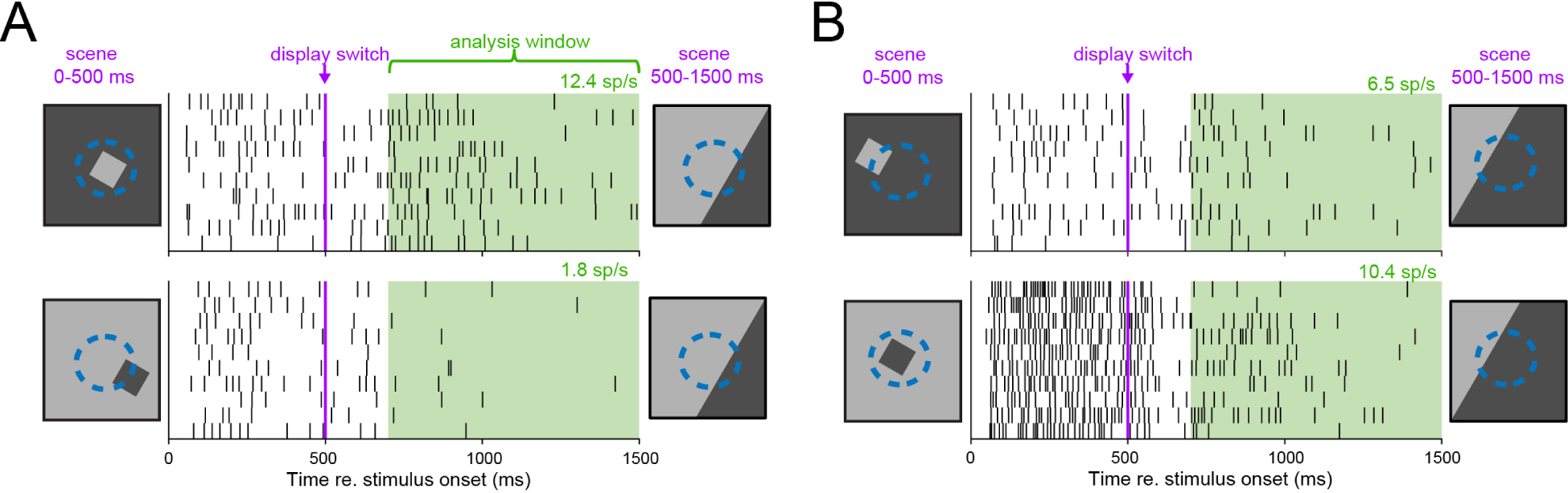
Dot rasters for the same unit as in Figure 2A,B, for stimulus sequences that have the opposite luminance contrast polarity as in Figure 2A,B but are otherwise identical.

**Figure S2.**
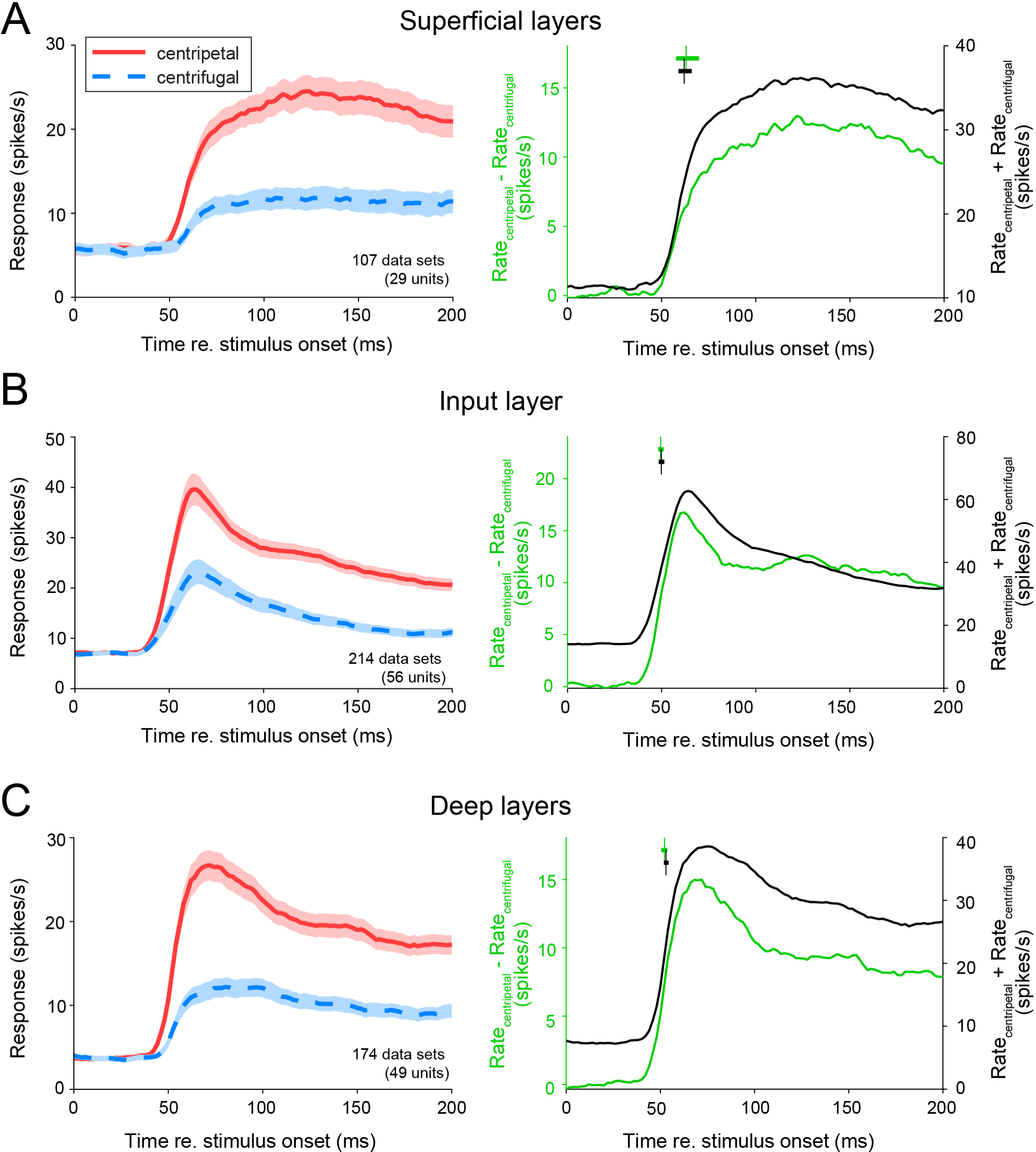
Onset of grouping signal computed separately for units in different laminar compartments. Similar as Figure 3A,B, for the subpopulations of grouping cells recorded respectively in superficial layers (A), granular (input) layer (B) and deep layers (C). Latency values in right panels: superficial layers difference function: 63.0 ms (95% CI [57.6 69.8] ms); superficial layers sum function: 62.0 ms (95% CI [59.0 66.0] ms); input layer difference function: 49.5 ms (95% CI [48.3 51.1] ms); input layer sum function: 49.9 ms (95% CI [48.5 51.4] ms); deep layers difference function: 52.2 ms (95% CI [50.5 53.7] ms); deep layers sum function: 53.0 ms (95% CI [51.8 54.3] ms).

**Figure S3.**
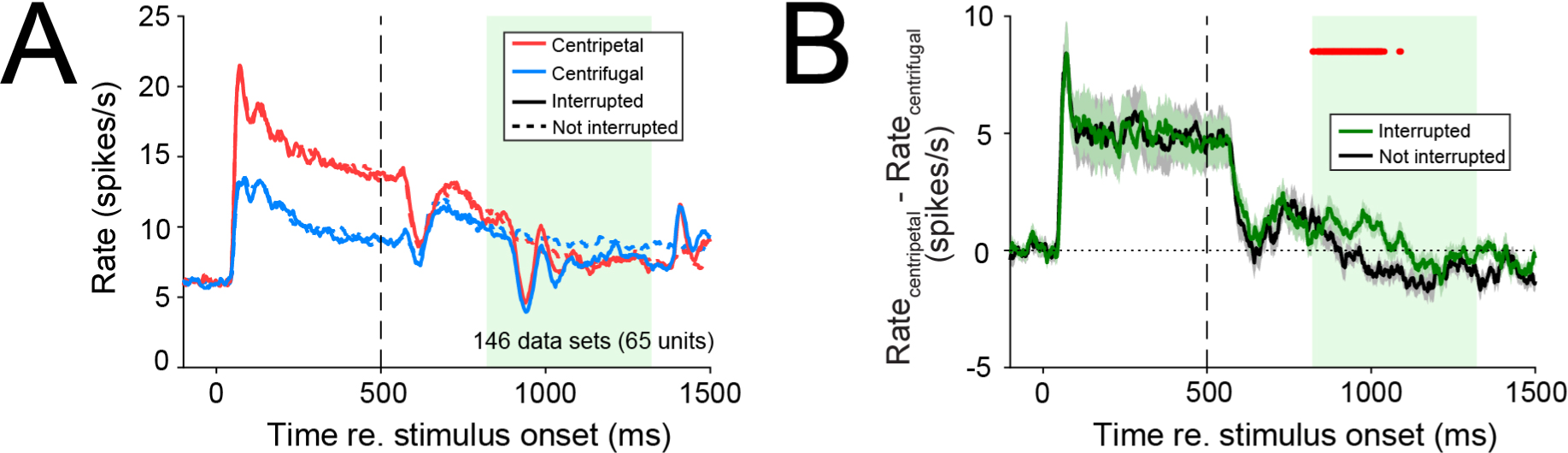
No persistent grouping signal during interrupt trials for units that do not pass the grouping cell criterion. (A) Similar as Figure 5B for the population of non-grouping cells (146 data sets from 65 non-grouping cells in animal D). (B) Similar as Figure 5C for non-grouping cells.

## REFERENCES

1. Cavanagh, P. (2011). Visual cognition. Vision Research 51, 1538–1551.

2. Peters, B., and Kriegeskorte, N. (2021). Capturing the objects of vision with neural networks. Nat Hum Behav 5, 1127–1144. 10.1038/s41562-021-01194-6.

3. Nakayama, K., He, Z.J., and Shimojo, S. (1995). Visual surface representation: a critical link between lower-level and higher-level vision. In An invitation to cognitive science. Visual cognition: An invitation to cognitive science (The MIT Press), pp. 1–70.

4. Zhou, H., Friedman, H.S., and von der Heydt, R. (2000). Coding of border ownership in monkey visual cortex. J. Neurosci. 20, 6594–6611.

5. Hesse, J.K., and Tsao, D.Y. (2016). Consistency of Border-Ownership Cells across Artificial Stimuli, Natural Stimuli, and Stimuli with Ambiguous Contours. J. Neurosci. 36, 11338– 11349. 10.1523/JNEUROSCI.1857-16.2016.

6. Williford, J.R., and von der Heydt, R. (2016). Figure-Ground Organization in Visual Cortex for Natural Scenes. eNeuro 3. 10.1523/ENEURO.0127-16.2016.

7. Franken, T.P., and Reynolds, J.H. (2021). Columnar processing of border ownership in primate visual cortex. Elife 10, e72573. 10.7554/eLife.72573.

8. Zhang, N.R., and von der Heydt, R. (2010). Analysis of the context integration mechanisms underlying figure-ground organization in the visual cortex. J Neurosci 30, 6482–6496. 10.1523/JNEUROSCI.5168-09.2010.

9. Craft, E., Schütze, H., Niebur, E., and von der Heydt, R. (2007). A neural model of figure-ground organization. J Neurophysiol 97, 4310–4326. 10.1152/jn.00203.2007.

10. Jehee, J.F.M., Lamme, V.A.F., and Roelfsema, P.R. (2007). Boundary assignment in a recurrent network architecture. Vision Res 47, 1153–1165. 10.1016/j.visres.2006.12.018.

11. Layton, O.W., and Yazdanbakhsh, A. (2015). A neural model of border-ownership from kinetic occlusion. Vision Res 106, 64–80. 10.1016/j.visres.2014.11.002.

12. Mehrani, P., and Tsotsos, J.K. (2021). Early recurrence enables figure border ownership. Vision Res 186, 23–33. 10.1016/j.visres.2021.04.009.

13. Zhu, S., Oh, Y., Trepka, E., Chen, X., and Moore, T. (2023). Border Ownership Selectivity and Laminar Connectivity in Single Columns of Macaque V1 Measured with High-Density Neuropixels Recordings. Society for Neuroscience Annual Meeting 2023 abstract. Poster presentation PSTR083.09 / W18. https://www.abstractsonline.com/pp8/#!/10892/presentation/26166

14. Douglas, R.J., and Martin, K.A.C. (2004). Neuronal circuits of the neocortex. Annu Rev Neurosci 27, 419–451. 10.1146/annurev.neuro.27.070203.144152.

15. Zarrinpar, A., and Callaway, E.M. (2016). Functional Local Input to Layer 5 Pyramidal Neurons in the Rat Visual Cortex. Cerebral Cortex 26, 991–1003. 10.1093/cercor/bhu268.

16. O’Herron, P., and von der Heydt, R. (2009). Short-term memory for figure-ground organization in the visual cortex. Neuron 61, 801–809. 10.1016/j.neuron.2009.01.014.

17. O’Herron, P., and von der Heydt, R. (2013). Remapping of border ownership in the visual cortex. J Neurosci 33, 1964–1974. 10.1523/JNEUROSCI.2797-12.2013.

18. Hu, B., von der Heydt, R., and Niebur, E. (2019). Figure-Ground Organization in Natural Scenes: Performance of a Recurrent Neural Model Compared with Neurons of Area V2. eNeuro 6, ENEURO.0479-18.2019. 10.1523/ENEURO.0479-18.2019.

19. Nakayama, K. (2005). Resolving border disputes in midlevel vision. Neuron 47, 5–8. 10.1016/j.neuron.2005.06.025.

20. von der Heydt, R. (2015). Figure-ground organization and the emergence of proto-objects in the visual cortex. Front Psychol 6, 1695. 10.3389/fpsyg.2015.01695.

21. Zhu, S.D., Zhang, L.A., and von der Heydt, R. (2020). Searching for object pointers in the visual cortex. J Neurophysiol 123, 1979–1994. 10.1152/jn.00112.2020.

22. Anderson, J.C., and Martin, K.A.C. (2006). Synaptic connection from cortical area V4 to V2 in macaque monkey. Journal of Comparative Neurology 495, 709–721. 10.1002/cne.20914.

23. Ungerleider, L.G., Galkin, T.W., Desimone, R., and Gattass, R. (2008). Cortical connections of area V4 in the macaque. Cereb Cortex 18, 477–499. 10.1093/cercor/bhm061.

24. Markov, N.T., Ercsey-Ravasz, M.M., Ribeiro Gomes, A.R., Lamy, C., Magrou, L., Vezoli, J., Misery, P., Falchier, A., Quilodran, R., Gariel, M.A., et al. (2014). A weighted and directed interareal connectivity matrix for macaque cerebral cortex. Cereb Cortex 24, 17–36. 10.1093/cercor/bhs270.

25. Pasupathy, A., Popovkina, D.V., and Kim, T. (2020). Visual Functions of Primate Area V4. Annu Rev Vis Sci 6, 363–385. 10.1146/annurev-vision-030320-041306.

26. Markov, N.T., Vezoli, J., Chameau, P., Falchier, A., Quilodran, R., Huissoud, C., Lamy, C., Misery, P., Giroud, P., Ullman, S., et al. (2014). Anatomy of hierarchy: feedforward and feedback pathways in macaque visual cortex. J Comp Neurol 522, 225–259. 10.1002/cne.23458.

27. Nandy, A.S., Nassi, J.J., and Reynolds, J.H. (2017). Laminar Organization of Attentional Modulation in Macaque Visual Area V4. Neuron 93, 235–246. 10.1016/j.neuron.2016.11.029.

28. Lu, Y., Yin, J., Chen, Z., Gong, H., Liu, Y., Qian, L., Li, X., Liu, R., Andolina, I.M., and Wang, W. (2018). Revealing Detail along the Visual Hierarchy: Neural Clustering Preserves Acuity from V1 to V4. Neuron 98, 417–428.e3. 10.1016/j.neuron.2018.03.009.

29. Pettine, W.W., Steinmetz, N.A., and Moore, T. (2019). Laminar segregation of sensory coding and behavioral readout in macaque V4. Proc Natl Acad Sci U S A 116, 14749– 14754. 10.1073/pnas.1819398116.

30. Westerberg, J.A., Schall, M.S., Maier, A., Woodman, G.F., and Schall, J.D. (2022). Laminar microcircuitry of visual cortex producing attention-associated electric fields. Elife 11, e72139. 10.7554/eLife.72139.

31. Davis, Z.W., Dotson, N.M., Franken, T.P., Muller, L., and Reynolds, J.H. (2023). Spike-phase coupling patterns reveal laminar identity in primate cortex. Elife 12, e84512. 10.7554/eLife.84512.

33. Sugihara, T., Qiu, F.T., and von der Heydt, R. (2011). The speed of context integration in the visual cortex. J Neurophysiol 106, 374–385. 10.1152/jn.00928.2010.

34. von der Heydt, R. (2023). Visual cortical processing—From image to object representation. Frontiers in Computer Science 5. 10.3389/fcomp.2023.1136987.

34. Zhaoping, L. (2005). Border ownership from intracortical interactions in visual area v2. Neuron 47, 143–153. 10.1016/j.neuron.2005.04.005.

36. Martin, A.B., and von der Heydt, R. (2015). Spike synchrony reveals emergence of proto-objects in visual cortex. J Neurosci 35, 6860–6870. 10.1523/JNEUROSCI.3590-14.2015.

36. Nandy, A.S., Sharpee, T.O., Reynolds, J.H., and Mitchell, J.F. (2013). The fine structure of shape tuning in area V4. Neuron 78, 1102–1115. 10.1016/j.neuron.2013.04.016.

38. Gallant, J.L., Connor, C.E., Rakshit, S., Lewis, J.W., and Van Essen, D.C. (1996). Neural responses to polar, hyperbolic, and Cartesian gratings in area V4 of the macaque monkey. J Neurophysiol 76, 2718–2739. 10.1152/jn.1996.76.4.2718.

38. Neupane, S., Guitton, D., and Pack, C.C. (2020). Perisaccadic remapping: What? How? Why? Rev Neurosci 31, 505–520. 10.1515/revneuro-2019-0097.

39. Golomb, J.D., and Mazer, J.A. (2021). Visual Remapping. Annu Rev Vis Sci 7, 257–277. 10.1146/annurev-vision-032321-100012.

40. Duhamel, J.R., Colby, C.L., and Goldberg, M.E. (1992). The updating of the representation of visual space in parietal cortex by intended eye movements. Science 255, 90–92. 10.1126/science.1553535.

41. Walker, M.F., Fitzgibbon, E.J., and Goldberg, M.E. (1995). Neurons in the monkey superior colliculus predict the visual result of impending saccadic eye movements. J Neurophysiol 73, 1988–2003. 10.1152/jn.1995.73.5.1988.

42. Umeno, M.M., and Goldberg, M.E. (2001). Spatial processing in the monkey frontal eye field. II. Memory responses. J Neurophysiol 86, 2344–2352. 10.1152/jn.2001.86.5.2344.

43. Churan, J., Guitton, D., and Pack, C.C. (2011). Context dependence of receptive field remapping in superior colliculus. J Neurophysiol 106, 1862–1874. 10.1152/jn.00288.2011.

44. Inaba, N., and Kawano, K. (2014). Neurons in cortical area MST remap the memory trace of visual motion across saccadic eye movements. Proc Natl Acad Sci U S A 111, 7825–7830. 10.1073/pnas.1401370111.

45. Nakamura, K., and Colby, C.L. (2002). Updating of the visual representation in monkey striate and extrastriate cortex during saccades. Proc Natl Acad Sci U S A 99, 4026–4031. 10.1073/pnas.052379899.

46. Neupane, S., Guitton, D., and Pack, C.C. (2016). Two distinct types of remapping in primate cortical area V4. Nat Commun 7, 10402. 10.1038/ncomms10402.

47. Marino, A.C., and Mazer, J.A. (2018). Saccades Trigger Predictive Updating of Attentional Topography in Area V4. Neuron 98, 429–438.e4. 10.1016/j.neuron.2018.03.020.

48. Olshausen, B.A., Anderson, C.H., and Essen, D.V. (1993). A neurobiological model of visual attention and invariant pattern recognition based on dynamic routing of information. J. Neurosci. 13, 4700–4719. 10.1523/JNEUROSCI.13-11-04700.1993.

49. Bushnell, B.N., Harding, P.J., Kosai, Y., and Pasupathy, A. (2011). Partial occlusion modulates contour-based shape encoding in primate area V4. J Neurosci 31, 4012–4024. 10.1523/JNEUROSCI.4766-10.2011.

50. Mysore, S.G., Vogels, R., Raiguel, S.E., and Orban, G.A. (2006). Processing of kinetic boundaries in macaque V4. J Neurophysiol 95, 1864–1880. 10.1152/jn.00627.2005.

51. Mysore, S.G., Vogels, R., Raiguel, S.E., and Orban, G.A. (2008). Shape selectivity for camouflage-breaking dynamic stimuli in dorsal V4 neurons. Cereb Cortex 18, 1429–1443. 10.1093/cercor/bhm176.

52. Anstis, S. (1970). Phi movement as a subtraction process. Vision research 10. 10.1016/0042-6989(70)90092-1.

53. Kolers, P.A., and Pomerantz, J.R. (1971). Figural change in apparent motion. J Exp Psychol 87, 99–108. 10.1037/h0030156.

55. Kolers, P.A., and von Grünau, M. (1976). Shape and color in apparent motion. Vision Research 16, 329–335. 10.1016/0042-6989(76)90192-9.

55. Ruiz, O., Lustig, B.R., Nassi, J.J., Cetin, A., Reynolds, J.H., Albright, T.D., Callaway, E.M., Stoner, G.R., and Roe, A.W. (2013). Optogenetics through windows on the brain in the nonhuman primate. J Neurophysiol 110, 1455–1467. 10.1152/jn.00153.2013.

56. Hwang, J., Mitz, A.R., and Murray, E.A. (2019). NIMH MonkeyLogic: Behavioral control and data acquisition in MATLAB. J Neurosci Methods 323, 13–21. 10.1016/j.jneumeth.2019.05.002.

57. Takeuchi, D., Hirabayashi, T., Tamura, K., and Miyashita, Y. (2011). Reversal of interlaminar signal between sensory and memory processing in monkey temporal cortex. Science 331, 1443–1447. 10.1126/science.1199967.

58. Yger, P., Spampinato, G.L., Esposito, E., Lefebvre, B., Deny, S., Gardella, C., Stimberg, M., Jetter, F., Zeck, G., Picaud, S., et al. (2018). A spike sorting toolbox for up to thousands of electrodes validated with ground truth recordings in vitro and in vivo. Elife 7, e34518. 10.7554/eLife.34518.

59. Keliris, G.A., Li, Q., Papanikolaou, A., Logothetis, N.K., and Smirnakis, S.M. (2019). Estimating average single-neuron visual receptive field sizes by fMRI. Proc Natl Acad Sci U S A 116, 6425–6434. 10.1073/pnas.1809612116.

60. Thompson, K.G., Hanes, D.P., Bichot, N.P., and Schall, J.D. (1996). Perceptual and motor processing stages identified in the activity of macaque frontal eye field neurons during visual search. J Neurophysiol 76, 4040–4055. 10.1152/jn.1996.76.6.4040.

